# Bee-associated fungi mediate effects of fungicides on bumble bees

**DOI:** 10.1101/2021.09.06.459164

**Authors:** Danielle Rutkowski, Eliza Litsey, Isabelle Maalouf, Rachel L. Vannette

**Affiliations:** Department of Entomology and Nematology, University of California Davis, Davis, CA 95616, United States; Department of Biology, University of Nevada Reno, Reno, NV, 89557

**Keywords:** *Ascosphaera*, bee microbiome, *Bombus*, propiconazole, *Starmerella*, yeast

## Abstract

Bumble bees are important pollinators that face threats from multiple sources, including agrochemical application. Declining bumble bee populations have been linked to fungicide application, which could directly affect the fungi often found in the stored food and GI tract of healthy bumble bees. Here, we test the hypothesis that fungicides impact bee health by disrupting bee-fungi interactions. We examine the interactive effects of the fungicide propiconazole and fungal supplementation on the survival, reproduction, and microbiome composition of microcolonies (queenless colonies) using two species, *Bombus vosnesenskii* and *B. impatiens*. We found that both bee species benefitted from fungi, but were differentially affected by fungicides. In *B. vosnesenskii*, fungicide exposure decreased survival while fungal supplementation mitigated fungicide effects. For *B. impatiens*, fungicide application had no effect, but fungal supplementation improved survival and offspring production. Fungicides reduced fungal abundance in *B. vosnesenskii* microcolonies, but not in *B. impatiens*, where instead fungal addition decreased fungal abundance. In *B. vosnesenskii*, the abundance of the pathogen *Ascosphaera* was negatively associated with survival, while the yeast *Zygosaccharomyces* was positively associated with survival. Our results highlight species-specific differences in response to fungicides and the nature of bee-fungi associations, and caution the use of results obtained using one species to predict responses of other species. These results demonstrate that fungicides can alter bee-fungi interactions with consequences for bee survival and reproduction, and suggest that exploring the mechanisms of such interactions, including interactions among fungi in the bee GI tract, may offer insights into bumble bee biology and conservation strategies.

**Significance statement:** Wild bumble bee populations are declining globally, and a major predictor of these declines is agricultural fungicide application. We test the hypothesis that bee-associated fungi mediate fungicide effects on bees, examining how fungicide exposure and subsequent fungal supplementation impact bumble bee survival and microbiome. Fungal supplementation enhanced survival in both bumble bee species tested here, with fungi mediating effects of fungicide exposure in one species. Differences in bee survival were associated with changes in the gut mycobiome: negatively with the pathogen *Ascosphaera* and positively with *Zygosaccharomyces* yeasts. This study highlights the importance of the mycobiome in bumble bee health both as a mechanism by which fungicides impact bumble bees and as an avenue of further research on beneficial bee-associated fungi.

## Introduction

Native bees, including bumble bees, are important pollinators of both native plants and agricultural crops [1,2]. Bumble bee (*Bombus* spp.) population decline has been well-documented [3,4], likely driven by multiple interacting factors including habitat degradation, pathogen prevalence, climate change, and agrochemical exposure [3,4,5,6]. On a landscape scale, the strongest predictor of bumble bee population decline is fungicide use [7], and experimental work shows that fungicide application can slow *B. impatiens* colony growth, resulting in smaller final colony size [8,9]. These studies demonstrate the need to understand the mechanisms through which fungicides impact bee health, and possible strategies to mitigate these effects.

The mechanisms behind fungicide effects on bumble bee colonies remain largely untested. One possible mechanism underlying fungicide impacts on bumble bee health is the disruption of bee-fungi associations. Bumble bees have a gut microbiome that contributes to bee nutrition and immune function [10,11]. Most research on bee microbiomes has focused on bacterial gut communities which are beneficial symbionts for bumble bees as well as other corbiculate bees [52]. However, bumble bees interact with fungi frequently and in multiple locations, although bee interactions with non-pathogenic fungi remain largely uncharacterized (but see [19]). Indeed, fungi, particularly yeasts, are commonly isolated from healthy bumble bees, including from the crop and gut [12,13,14], and from pollen provisions and stored nectar within the nest [15,16]. Bumble bees also exhibit behavioral preferences for yeast-containing flowers, preferentially foraging on flowers containing yeasts over flowers without yeasts [17,18]. Some evidence indicates that removal of fungi from food stores of bumble bees decreases the survival of developing larvae [15,19], and addition of certain yeast species to *B. terrestris* diets increases colony growth while also slowing the growth of the bumble bee pathogen *Crithidia bombi* [19]. Fungicides reduce the growth of nectar- and bee-associated yeasts in culture [9,15,20,21], and thus one clear, but untested, route by which fungicides could impact bumble bees is disruption of their associations with endosymbiotic fungi.

In this study, we examined effects of fungicide application and fungal addition on worker bumble bee health and microbiome composition. We tested the hypotheses that (1) fungicide exposure affects bumble bee health, that (2) supplementation with bumble bee-associated fungi following fungicide exposure can mitigate the effects of fungicide, and that (3) changes in the composition and abundance of fungi within the gut microbiome mediate fungicide effects. To test these hypotheses, we used two species of bumble bee, the yellow-faced bumble bee, *Bombus vosnesenskii*, and the common eastern bumble bee, *B. impatiens*. We tested the effects of propiconazole, a widely used agricultural fungicide, and subsequent addition of fungi isolated from each bee species, and examined effects on feeding, foraging behavior, offspring production, survival, and microbiome composition.

## Results

### Microcolony survival

Fungicide and fungi addition affected microcolony survival in *B. vosnesenskii* and *B. impatiens*, but bee species differed in their responses. For *B. vosnesenskii*, fungicide without fungi addition reduced microcolony survival, with only about 50% of colonies surviving to day 20. However, microcolonies given fungi following fungicide exposure were indistinguishable from control microcolonies, both with nearly 85% survival to day 20 (fungicide x fungi treatments, χ^2^ = 5.55, df = 1, p = 0.02, Fig 1A). In contrast, *B. impatiens* microcolony survival did not change with fungicide application (χ^2^ = 0.049, p = 0.43), but instead greatly increased with fungal addition (χ^2^ = 21, df = 1, p < 0.001, Fig 1B). Between 85-100% of microcolonies given fungi survived to day 30, while only 40% of microcolonies that did not receive fungi survived to that point, regardless of fungicide application.

**Figure 1.**
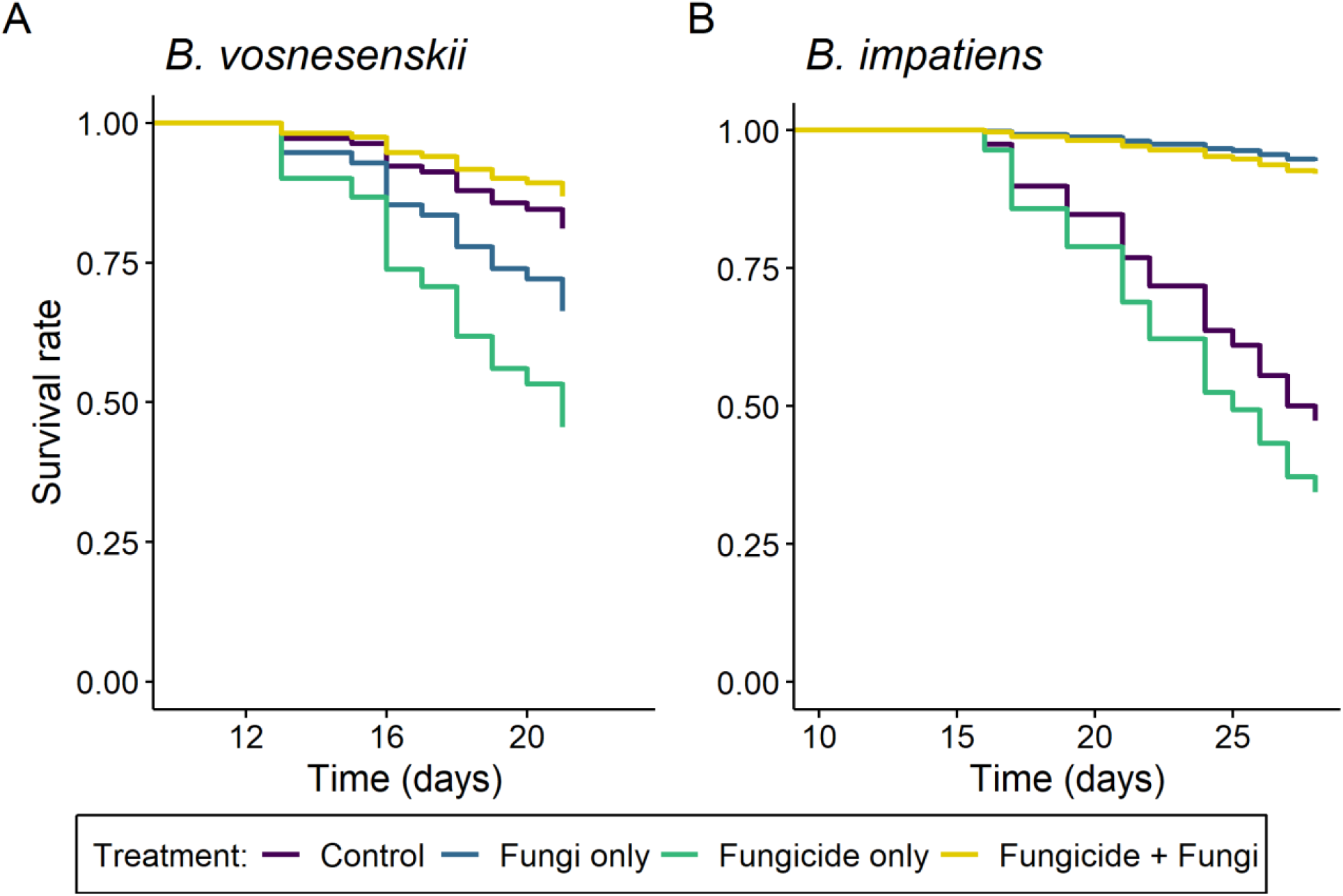
Survival over time of microcolonies of (A) *B. vosnesenskii* and (B) *B. impatiens* in different fungicide and fungi treatment groups. Survival of *B. vosnesenskii* microcolonies was treatment-dependent (χ^2^ = 8.08, df = 3, p = 0.04), with microcolonies given fungicide only experiencing the highest mortality. Treatment also impacted *B. impatiens* microcolony survival (χ ^2^ = 20, df = 3, p < 0.001), with microcolonies receiving fungal supplementation surviving longer, while fungicide treatment had no effect. N = 15 microcolonies per treatment per species.

### Nectar consumption

For both bee species, fungi but not fungicide affected nectar feeding. Fungal addition increased *B. vosnesenskii* per-bee nectar consumption by 12% (F_1,48_ = 4.43, p = 0.040), while neither previous fungicide treatment (F_1,46_ = 0.12, p = 0.74) nor the interaction between fungicide treatment and fungi treatment (F_1, 51_ = 1.23, p = 0.27) impacted consumption. In *B. impatiens*, fungal addition increased per-bee nectar consumption by 24% compared to control nectar (F_1,55_ = 20.44, p < 0.001). Previous fungicide treatment (F_1,55_ = 2.55, p = 0.12) and the interaction between fungicide treatment and fungi treatment (F_1, 55_ = 1.73, p = 0.19) had no effect on *B. impatiens* nectar consumption. Fungicide addition did not affect nectar consumption by either bee species (*B. vosnesenskii*, F_1,51_ = 0.10, p = 0.75; *B. impatiens*, F_1,58_ = 0.0038, p = 0.95).

### Y-tube preference trials

Full y-tube results are contained in Supplemental Results. Briefly, *B. vosnesenskii* workers were tested in two preference trials, one between fungicide and control nectar volatiles and one between yeast (*Metschnikowia reukaufii*) and control nectar volatiles. Workers showed no preference between fungicide and control volatiles, but preferred control volatiles over yeasts.

### Offspring production

Offspring production was not quantified for *B. vosnesenskii* as no offspring were produced. In *B. impatiens* microcolonies, fungal addition increased egg abundance (hurdle model, p = 0.007, Fig 2A), while neither fungicide treatment (p = 0.90) nor the fungicide x fungi interaction affected egg abundance (p = 0.82). Fungal addition also increased egg mass (F_1,18_ = 5.02, p = 0.038, Fig 2B), but egg mass was not affected by fungicide treatment (F_1,18_ = 0.59, p = 0.59) nor the fungicide x fungi interaction (F_1,18_ = 0.026, p = 0.87).

**Figure 2.**
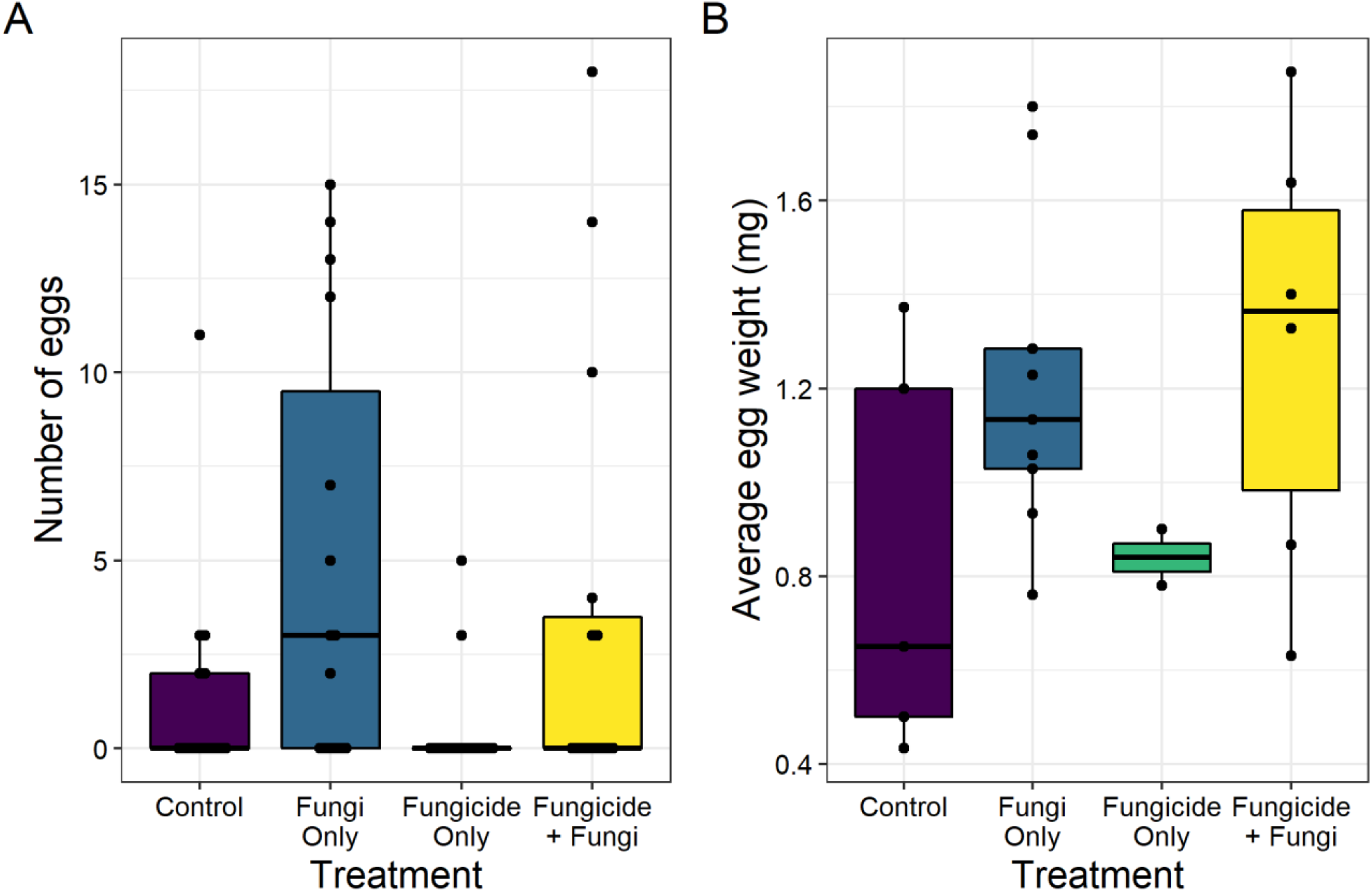
The (A) number and (B) average weight of eggs laid in microcolonies of *B. impatiens* across fungicide and fungi treatments. Both egg abundance (p = 0.006) and egg weight (p = 0.038) were positively affected by fungal supplementation, regardless of fungicide treatment. N = 15 microcolonies per treatment.

## Fungal microbiome

### Fungal abundance (qPCR)

To examine if fungicide or fungi addition affected total fungal abundance, we quantified ITS copy number using qPCR. In *B. vosnesenskii*, fungicide application without subsequent fungal treatment reduced ITS copy number compared to all other treatments (F_1,58_ = 4.48, p = 0.04, Fig 3A). In addition, fungal copy number was greater in the gut than in the crop (F_1,53_ = 12.58, p < 0.001), and there were no interactions between treatments and organ (p > 0.50 for all comparisons). A contrasting response in ITS copy number was found in B. *impatiens* microcolonies, where fungi-treated microcolonies had lower ITS copy number than microcolonies not treated with fungi (F_1,58_ = 8.76, p = 0.004, Fig 3B). ITS copy number was not affected by fungicide treatment (F_1,58_ = 1.89, p = 0.17) nor the fungicide x fungi interaction (F_1,58_= 0.87, p = 0.35). Gut samples contained higher copy number than the crops, though this difference was only marginally significant (F_1,58_ = 3.84, p = 0.055), and there were no interactions between treatments and organ (p > 0.50 for all comparisons).

**Figure 3.**
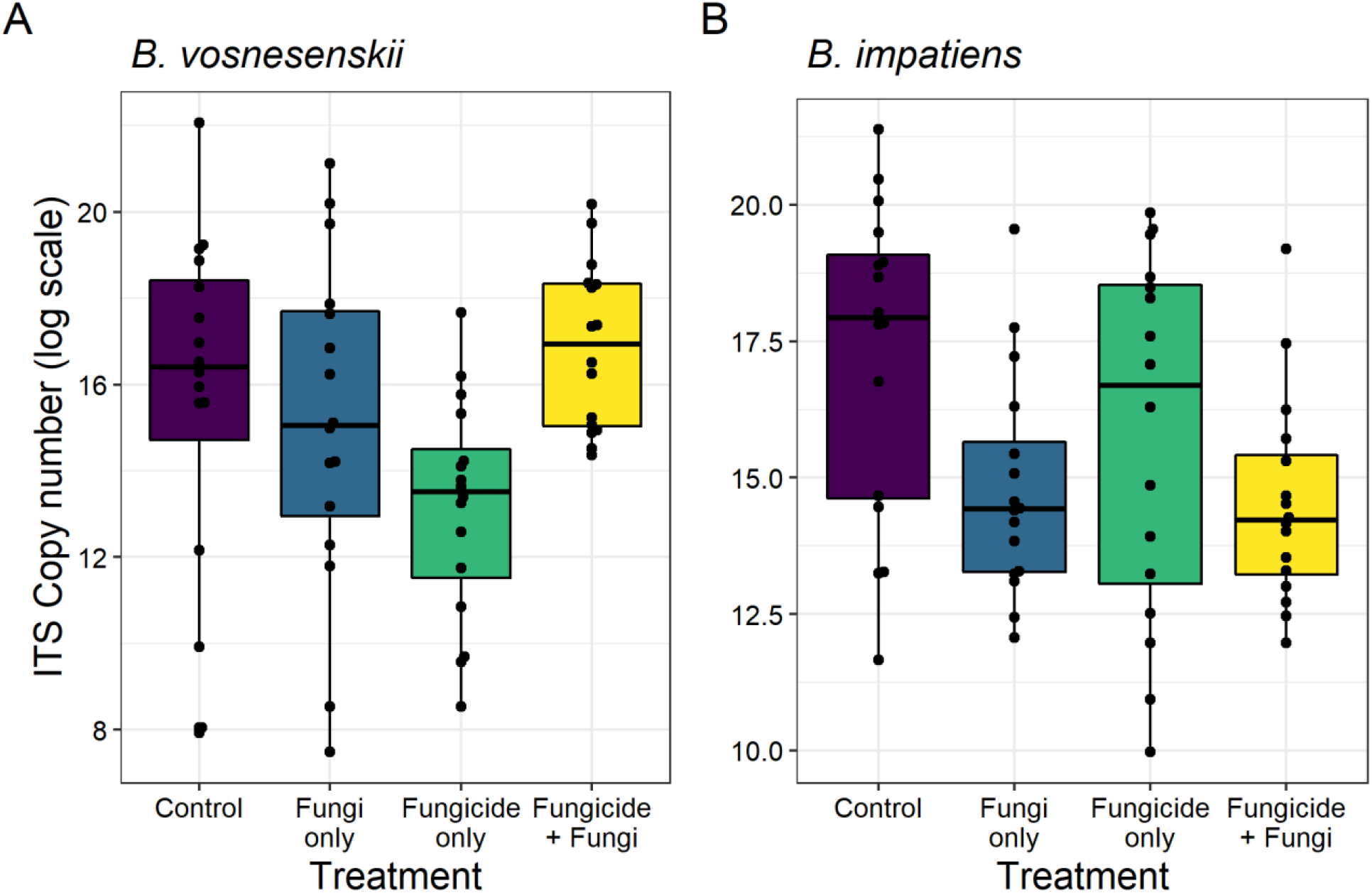
Fungal copy number in A) *B. vosnesenskii* and B) *B. impatiens* microcolonies by treatment. *B. vosnesenskii* fungal abundance depended on both fungicide and fungi treatments (F_1,58_ = 4.48, p = 0.04), while *B. impatiens* fungal abundance was only affected by fungi treatment (F_1,58_ = 8.76, p = 0.004). N = 8 microcolonies per treatment per species.

### Amplicon sequencing

Following quality filtering and preprocessing steps, a total of 2,169,875 ITS sequences were obtained from 64 *B. vosnesenskii* samples, ranging from 208 to 118,191 per sample (33,904 ± 529, mean ± SE) which clustered into a total of 349 ASVs. All samples were saturating (Figure S1A). Ascomycete fungi accounted for 96% of sequences, comprised of families Saccharomycetaceae (36%), Debaryomycetaceae (22%), Ascosphaeraceae (16%), and Aspergillaceae (11%), among others.

A total of 598,998 ITS sequences were obtained from 64 *B. impatiens* samples, ranging from 8 to 65,969 per sample (9,359 ± 208). Sequences were clustered into 54 ASVs. All sampling curves were saturating (Figure S1B). Ascomycetes accounted for 99.6% of sequences, and families included Saccharomycetaceae (66%), Ascosphaeraceae (12%), Aureobasidiaceae (10%), with the remainder comprised of Debaryomycetaceae and Aspergillaceae.

### Alpha diversity

We examined if treatments affected alpha diversity of the fungal microbiome using Shannon diversity indices. In *B. vosnesenskii*, Shannon diversity was not affected by fungicide treatment (F_1,75_ = 0.15, p = 0.70), fungi treatment (F_1,59_ = 0.041, p = 0.84), nor their interaction (F_1,55_ = 0.002, p = 0.96, Fig 4A). In contrast, for *B. impatiens* microcolonies, treatments interacted to affect Shannon diversity (F_1,59_ = 4.47, p = 0.039, Fig 4B), where fungi addition increased Shannon diversity, especially in microcolonies that had previously received fungicide. Fungal communities from crop samples were more diverse than gut samples for *B. impatiens* (F_1,59_ = 12.02, p < 0.001), but no difference was detected for *B. vosnesenskii* (F_1,55_ = 1.64, p = 0.21).

**Figure 4.**
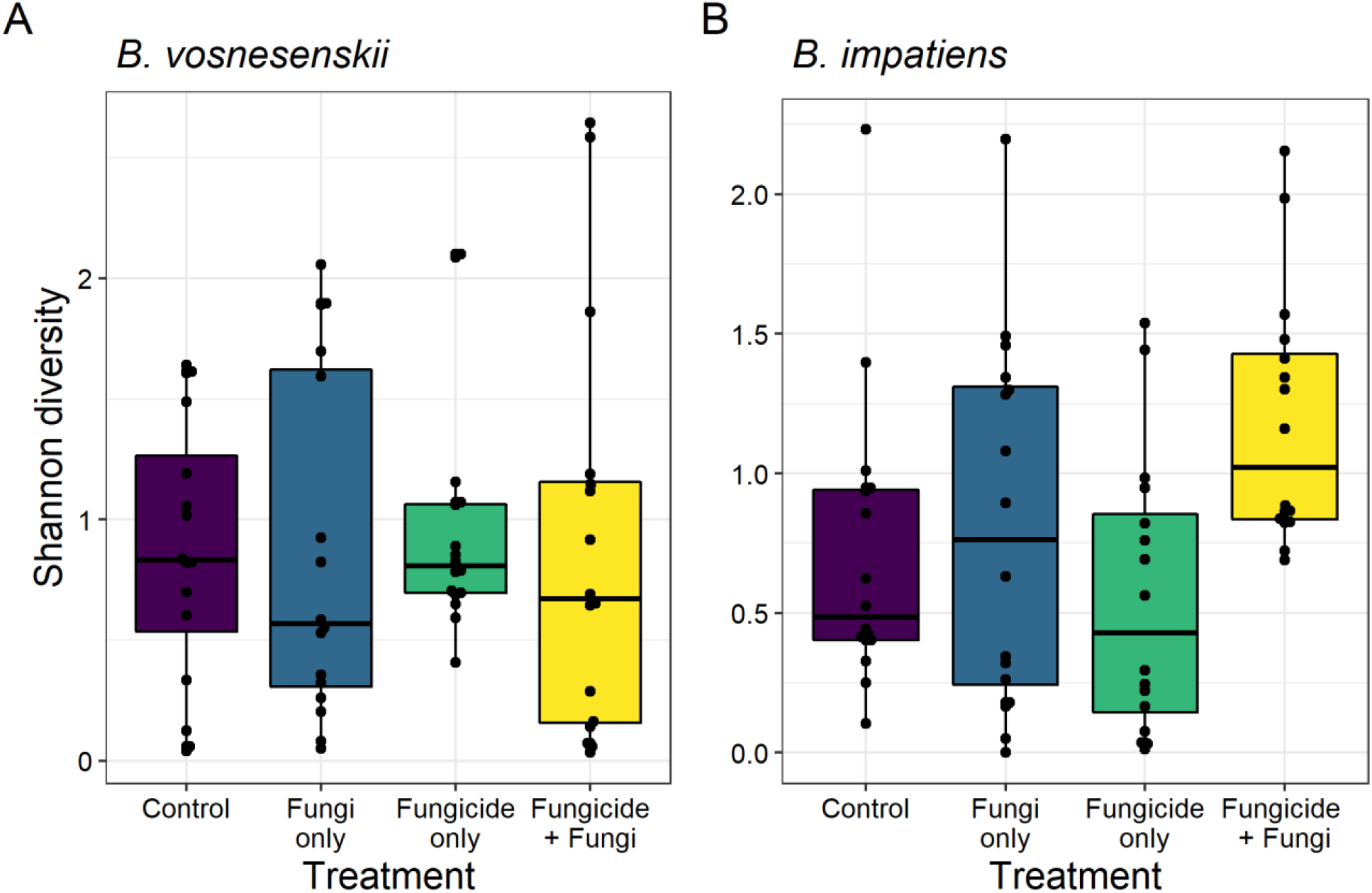
Shannon diversity of A) *B. vosnesenskii* and B) *B. impatiens* fungal microbiome. In *B. vosnesenskii*, Shannon diversity was unaffected by treatment (F_1,55_ = 0.002, p = 0.96), but in *B. impatiens* diversity was impacted by the interaction of fungicide and fungi treatment (F_1,59_ = 4.47, p = 0.039). N = 8 microcolonies per treatment per species.

### Fungal composition

Next, we examined if fungal community composition varied with treatments or was associated with microcolony survival. For *B. vosnesenskii*, fungi treatment (PERMANOVA, F_1,18_ = 2.78, R^2^ = 0.038, p = 0.002) and the interaction between fungicide treatment and fungi treatment (F_1,18_ = 3.99, R^2^ = 0.054, p = 0.001, Fig 5A, Fig S2A) affected fungal community composition, while fungicide treatment alone had no effect (F_1,18_ = 1.54, R^2^ = 0.021, p = 0.088). Crop and gut samples did not differ in community composition (F_1,18_ = 0.57, p = 0.92). Notably, *B. vosnesenskii* source colonies differed in their fungal community compositions (F_7,18_ = 2.26, R^2^ = 0.21, p = 0.001), and microcolonies sourced from different colonies differed in fungal community response to fungicide and fungi treatment (F_1,18_ = 2.46, R^2^ = 0.033, p = 0.003). Among-sample variance did not differ between fungicide treatments (betadisper, F_1,62_ = 0.30, p = 0.58), fungi treatments (F_1,62_ = 0.33, p = 0.56), or crop and gut samples (F_1,62_ = 0.14, p = 0.71), but was different among source colonies (F_7,56_ = 3.74, p = 0.004).

**Figure 5.**
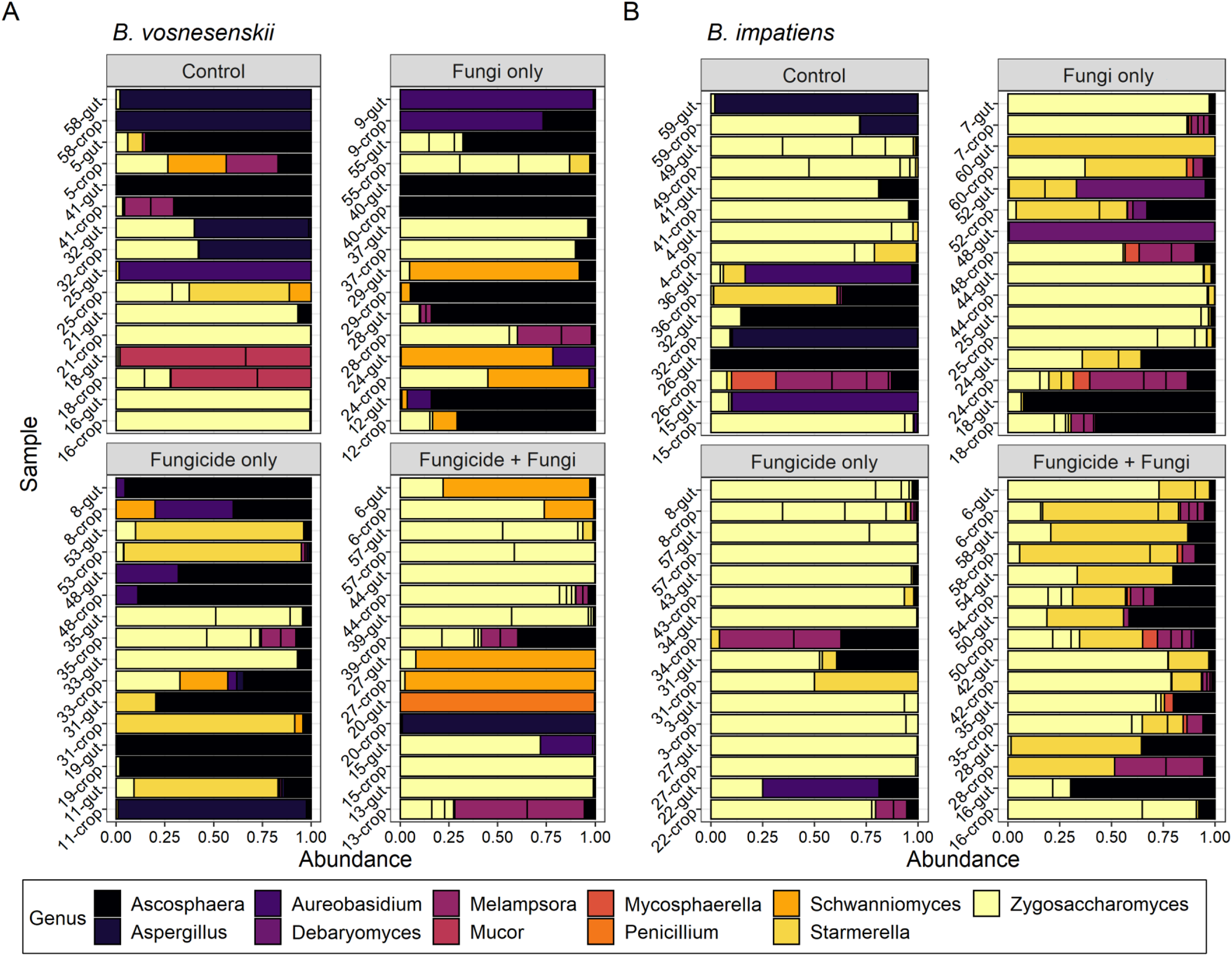
Fungal community composition of (A) *B. vosnesenskii* and (B) *B. impatiens* samples across fungicide and fungi treatments. The relative abundance of the top 20 taxa are shown are shown for each sample, and different colors correspond to different fungal genera. **Supplemental materials for: Bee-associated fungi mediate effects of fungicides on bumble bees**

In *B. impatiens*, fungal species composition differed between fungi treatments (F_1,58_ = 3.79, R^2^ = 0.056, p = 0.006, Fig 5B, Fig S2B), but not between fungicide treatments (F_1,58_ = 1.31, p = 0.25) or their interaction (F_1,58_ = 1.24, p = 0.24). Communities were similar between crop and gut samples (F_1,58_ = 1.77, p = 0.11) and between source colonies (F_1,58_ = 1.23, p = 0.27). Variance was homogeneous across fungicide treatments (F_1,62_ = 0.43, p = 0.55), fungi treatments (F_1,62_ = 0.024, p = 0.87), crop and gut samples (F_1,62_ = 0.44, p = 0.51), and source colonies (F_1, 62_ = 0.059, p = 0.80).

To examine if mycobiome composition (based on relative abundances) was associated with microcolony survival, we ran a Cox proportional hazards analysis, with fungal PCoA axes as predictors for each species separately. For *B. vosnesenskii*, PCoA axis 2 was significantly associated with survival (χ^2^ = 4.60, df = 1, p = 0.032), but axis 1 was not (χ^2^ = 2.89, df =1, p = 0.089). For *B. impatiens*, axis 2 was marginally associated with variation in survival (χ^2^ = 3.69, df =1, p = 0.055), while axis 1 was not (χ^2^ = 0.37, df = 1, p = 0.54). See Supplemental Results for more details and differential abundance analyses.

### Abundance-corrected analyses

We hypothesized that the abundance of specific fungal groups may mediate the effects of fungicide or fungal addition observed above, including yeasts added to bee diets (*Starmerella, Zygosaccharomyces, Debaryomyces*) or the fungal pathogen *Ascosphaera*, which is often found in commercial pollen diets [32,34]. We used fungal copy number and relative abundance in the amplicon dataset to estimate total abundance of these four focal genera and examined their association with microcolony survival.

In *B. vosnesenskii*, fungicide-only treatment reduced *Zygosaccharomyces* abundance compared to all other treatments (F_1,56_ = 11.33, p = 0.001, Fig S4A). In addition, *Zygosaccharomyces* abundance was positively associated with microcolony survival (χ ^2^ = 4.57, df = 1, p = 0.033). Abundance of the putatively beneficial yeast *Starmerella* was not affected by treatments (p > 0.06 for all comparisons, Fig S4B) and was not associated with differences in survival (χ ^2^ = 0.15, df = 1, p = 0.7). Abundance of the pathogen *Ascosphaera* was low in control microcolonies and those receiving fungicide and highest in the fungi-only treatment (F_1,54_ = 14.00, p < 0.001, Fig S4C). Increasing *Ascosphaera* abundance was associated with reduced microcolony survival (χ ^2^ = 7.18, df = 1, p = 0.007).

In *B. impatiens*, similar fungal genera dominated the community as in *B. vosnesenskii*, but genera were differently affected by treatments. *Zygosaccharomyces* abundance was unaffected by fungicide treatment (F_1,55_ = 0.03, p = 0.86) or the interaction between treatments (F_1,55_ = 0.001, p = 0.97), but was reduced by fungal addition (F_1, 55_ = 6.79, p = 0.012, Fig S4A), despite being present in the inoculum provided to microcolonies. Interestingly, *Zygosaccharomyces* abundance was negatively correlated with microcolony survival, though only marginally (χ ^2^ = 3.22, df = 1, p = 0.07). *Starmerella* abundance was also impacted by the interaction of fungicide and fungi treatments (F_1,55_ = 5.12, p = 0.028, Fig S4B), and was lowest in the fungicide-only treatment. *Starmerella* abundance was not associated with survival (χ ^2^ = 0.01, df = 1, p = 0.9). *Debaryomyces* abundance was unaffected by fungicide treatment, fungi treatment, or their interaction (p > 0.10 for all comparisons) and was not associated with survival (χ ^2^ = 0.05, df = 1, p = 0.8). *Ascosphaera* abundance was reduced by fungicide (F_1,56_ = 7.35, p = 0.009, Fig S4C), but was relatively abundant in all other treatments, and was not associated with differences in survival (χ ^2^ = 0.24, df = 1, p = 0.6).

## Bacterial microbiome

Full bacterial microbiome results are contained in Supplemental Results. Briefly, in *B. vosnesenskii* microcolonies, bacterial alpha diversity was unaffected by fungicide and fungal treatment, but community composition was affected by the interaction of fungicide and fungi treatments (F_1,18_ = 2.46, R^2^ = 0.041, p = 0.019, Fig S4). In *B. impatiens* microcolonies, bacterial alpha diversity was lower in fungicide-treated microcolonies, but only in crop samples (F_1,50_ = 5.97, p = 0.018). Community composition was unaffected by fungicide treatment or fungi treatment.

## Discussion

Here, we show that fungi increase bumble bee microcolony survival and reproduction and that the effects of fungicides on bumble bees can be mediated by the fungal microbiome. Additionally, the two bumble bee species tested here showed qualitative differences in their responses to fungicide and fungi addition. In *Bombus vosnesenskii*, fungicide application reduced survival of microcolonies, but survival was restored to high levels when fungi were provided to bees following exposure. In *B. impatiens*, microcolonies were unaffected by fungicide application, but benefited from fungi supplementation. Distinct responses of bee species and their microbiomes to treatments shed light on the potential mechanisms of fungicide effects on bumble bees and bee-fungi interactions.

Our study provides strong evidence for negative effects of the widely used fungicide propiconazole on *B. vosnesenskii* survival, mediated by its effect on fungal abundance and composition in the bee gut. Although fungicide application has been shown to be a strong landscape-scale predictor of bumble bee decline in recent studies [7], short-term toxicity studies have shown relatively minimal effects of fungicides on bumble bees and honey bees [53,54], leading to the conclusion that fungicides pose minimal harm to pollinators. However, in longer-term studies, consumption of the azole propiconazole used here has been shown to cause both lethal [23,24] and sublethal effects on bumble bees such as decreased nectar consumption and reduced cell number in nests [22]. In these studies, a minimum of 2-3 weeks was required to detect negative effects of fungicides on survival and reproduction, suggesting that short-term toxicity studies can overlook delayed or indirect effects, similar to those we detected on bees and their associated fungi. Such effects may be more common among bees than is currently recognized [54]. However, we expect fungicide effects to vary among bee species as shown here, and among fungicide classes and formulations. Propiconazole is known to be highly toxic to bee- and nectar-associated fungi [20,21] and our study documented significant effects on fungal abundance and composition in one bee species. For *B. vosnesenskii*, propiconazole reduced the abundance of fungi including *Zygosaccharomyces*, but fungal reintroduction decreased fungicide-induced mortality. However, in *B. impatiens*, propiconazole had no detectable effect on fungal abundance nor composition. Bee species’ divergent responses to fungicide and fungi addition could be due to differences in initial microbiome composition and sensitivity to fungicides, or the nature of the isolated and re-introduced fungi. Each of these options could be examined in more detail in future experiments. Based on previous studies of fungicide effects on yeasts, we predict that azole fungicides may be more detrimental than other fungicide classes to bee-associated yeasts [20], but this remains to be experimentally examined.

Like fungicide application, the effects of fungal supplementation differed between species, and their divergent responses suggest that a few distinct mechanisms may mediate bee response to non-pathogenic fungi. In *B. vosnesenskii*, fungal treatment increased fungal abundance as expected, and although the reconstituted microbiome following fungal supplementation did not exactly mirror the initial community (Fig 5A), both were dominated by *Zygosaccharomyces* and low in *Ascosphaera*. In contrast, fungal addition to *B. impatiens* reduced overall fungal abundance, reducing *Zygosaccharomyces* abundance while *Starmerella* yeasts and *Ascosphaera* remained abundant.

Previous work has suggested that fungi in the diet can serve directly as food [25,26], that fungal metabolism may produce nutrients or important metabolites [27,33], or that beneficial fungi could suppress the growth of pathogens [19]. Fungi also have the potential to impact bee health through changes in behavior. Nectar yeasts can attract bumble bees to flowers [28,29], and here, yeasts stimulated feeding in both species. While our work cannot directly address the mechanism of fungal effect, we note that different mechanisms are implicated for the two bumble bee species used here. Additionally, fungal identity, not just abundance, seems to play an important role in fungal effects.

A closer examination of mycobiome responses in each species gives clues about potential mechanisms underlying bee response. In *B. vosnesenskii, Zygosaccharomyces* was highly abundant and was positively correlated with microcolony survival. *Zygosaccharomyces* is a common bee associate [30,31,32] that produces sterols that are vital for development in some bee species [27] but its metabolism when associated with bumble bees has not yet been examined. We also examined if correlations among taxa indicated the potential for pathogen suppression. Here, *Zygosaccharomyces* abundance was not correlated with *Ascosphaera* abundance (Supplemental Results), so we interpret these data as supporting a nutritional rather than pathogen suppression hypothesis, but both will require experimental validation. In contrast, the yeast nutrition hypothesis is not supported in *B. impatiens*. Instead, microcolonies performed most poorly when fungal abundance was greatest (and diversity lowest). Additionally, none of the focal fungal taxa were associated positively with survival. Therefore, it is possible that fungal addition improved *B. impatiens* health through other interactions that were not measured in this study, such as interaction with unmeasured bumble bee pathogens and parasites, although further studies would need to be performed to determine these mechanisms. In contrast to previous studies on *B. terrestris* [19], we did not detect a positive effect of *Starmerella* abundance on bumble bee survival in either species. It is possible that yeasts within this genus could have context-specific effects on bumble bees, or that different bee species benefit from specific species or strains of fungi.

Notably, bumble bee species also differed by a large margin in the diversity and composition of their fungal microbiome. Fungal communities in *B. vosnesenskii* hosted 349 fungal ASVs (mean 15 ASVs/sample), which was much more diverse than communities in *B. impatiens*, which hosted only 54 ASVs (mean 10 ASVs/sample). Moreover, wild-caught *B. vosnesenskii* queens produced colonies that differed significantly in mycobiome composition, explaining 21% of variation among samples. The commercially-reared *B. impatiens* colonies, in contrast, did not differ in fungal microbiome composition. We hypothesize that the process of commercial rearing may change fungal community composition and reduce fungal diversity [19]. Other studies have observed higher bacterial diversity in the GI tract of wild bumble bees compared to commercial bumble bees [11], supporting this hypothesis. Not only the diversity, but also the identity and functions of the microbiome differed between bee species and could be a product of rearing conditions. For example, the abundance of the bee pathogen *Ascosphaera* was negatively correlated with survival only in *B. vosnesenskii*, despite being prevalent at high levels in both species. *Ascosphaera* is commonly found in honey bee pollen used for commercial bumble bee rearing [32,34], so routine exposure to *Ascosphaera* may generate selection for reduced susceptibility in commercial species like *B. impatiens*, as has been found in a previous studies comparing the susceptibility of wild and commercial *B. terrestris* to *Crithidia bombi* [11].

## Conclusion*s*

The widespread use of fungicides particularly during crop blooms can spill over to affect managed and native pollinators. The results presented here suggest that fungi associated with bumble bees mediate effects of fungicides on bees, and fungal reintroduction could potentially mitigate harm following exposure. Future studies will be required to further characterize the mechanisms underlying bumble bee-yeast interactions and whether the effects of fungicide observed here are also observed in field conditions. Nevertheless, the results we describe here suggest that fungi can be influential members of the bee microbiome, and an underappreciated route through which agrochemicals harm pollinator populations.

## Methods

### Experimental overview

Bumble bee microcolonies were created either from colonies started with wild caught queens (*B. vosnesenskii*) or from purchased colonies (*B. impatiens*, Koppert, Howell MI). For each species, microcolonies were created and treated with fungicides and fungi in a factorial fashion, then survival and reproductive parameters were quantified. Finally, bees were dissected and gut contents extracted for microbiome analysis. Rearing and experimental details are described for each species below.

### Bombus vosnesenskii *rearing conditions*

In March 2019, emerging queens of *Bombus vosnesenskii*, an abundant bumble bee in the western United States [35], were collected from Monterey, CA. Captured *B. vosnesenskii* queens were transported to the Harry H. Laidlaw Jr. Honey Bee Research Facility at the University of California Davis. They were placed into rearing boxes (BioBest, USA) and each was provided with a pollen ball and BioGluc solution (BioBest, USA) as a carbohydrate source. Pollen balls were created by mixing together equal parts finely ground honeybee-collected pollen (Koppert, USA) and BioGluc solution. Queens were kept in a dark room at 27°C and 60% relative humidity. Pollen balls and BioGluc were replaced every 3-5 days as needed. We reared a total of nine source colonies for microcolony creation.

Newly-emerged workers were removed from *B. vosnesenskii* source colonies to create microcolonies. Each microcolony contained three workers from the same source colony, and was provided with a pollen ball. Microcolonies were assigned to one of two treatment groups: fungicide or control (n = 30/treatment). Control microcolonies received artificial nectar consisting of 60g sucrose, 120g glucose, 120g fructose, and 1g peptone in 950mL water. This nectar was autoclaved for 30 minutes prior to use, after which 50mL MEM non-essential amino acids (Corning, USA) was added to simulate the composition of natural floral nectar. Fungicide-treated microcolonies were given artificial nectar containing 7.5 ppm propiconazole (Quali-Pro, USA). Microcolonies fed on these nectar reservoirs for one week, and rate of consumption was recorded by massing nectar reservoirs before giving them to microcolonies and then again after five days. Following fungicide application, microcolony treatment groups received fungal treatments. Half of the microcolonies from each fungicide treatment group received sterile artificial nectar, and the other half was given nectar inoculated with a collection of fungi previously isolated from the crops of overwintering *B. vosnesenskii* (1×10^4^ cells/mL *Penicillium cyclopium, Starmerella bombi*, and *Zygosaccharomyces rouxii*), creating a total of four treatment groups (n = 15 microcolonies/treatment). Nectar consumption was again measured during the first week that microcolonies were placed on fungal treatments. Microcolonies were kept on their diet containing fungi or control nectar for two weeks total. Worker survival in each microcolony was measured daily, and microcolonies were terminated three weeks after their initiation. No microcolonies produced offspring during the duration of this experiment.

### Y tube assays

Foraging preferences of *B. vosnesenskii* workers from each microcolony were tested in two Y-tube assays. The first of these assays tested worker preference between volatiles of sterile artificial nectar and artificial nectar with added fungicide. The second assay tested worker preference between volatiles of sterile artificial nectar and artificial nectar inoculated with a common nectar yeast, *Metschnikowia reukaufii* (Supplemental Methods).

### Bombus impatiens *rearing conditions*

*B. impatiens* is a common bumble bee on the east coast of the United States that is commercially reared and used as an agricultural pollinator. This species is also commonly used in laboratory studies of bumble bee health and behavior. Colonies for this study were obtained from Koppert Biological Systems (Howell, MI, USA). The *B. vosnesenskii* microcolony protocol was repeated in fall 2019 using *Bombus impatiens*, with a few modifications. All microcolonies were created from two source colonies. Each microcolony contained five workers, and in addition to pollen balls and nectar, each microcolony was also provided with wax pellets (Beesworks organic yellow beeswax pellets) to encourage reproduction. Microcolonies were kept in incubators kept at 27°C and 60% relative humidity. Fungi-supplemented microcolonies received fungi previously isolated from the guts of *B. impatiens* workers (1×10^4^ cells/mL *Debaryomyces hansenii, Starmerella sorbosivorans*, and *Zygosaccharomyces rouxii*). Microcolonies were also kept on their fungi treatment diet for an extra week to allow offspring development, resulting in a total experiment time of four weeks. When microcolonies were terminated, each was dissected to count and weigh offspring (eggs, larvae, and pupae) present in each colony. Due to low abundance of larvae and pupae, only egg abundance and mass were compared between treatments. Preference assays were not performed for *B. impatiens* (but see [51] for similar assays).

### Dissection, DNA extraction, qPCR, and bioinformatics

Workers from all microcolonies were frozen at -20°C until dissection. Three workers from randomly chosen microcolonies of each treatment (n = 8 microcolonies/treatment/species) were dissected for DNA extraction. From each bee, the crop and midgut and hindgut, hereafter ‘gut’, were dissected and processed separately. To ensure each sample had sufficient DNA, organs from three bees of each microcolony were pooled to create a total of one crop and one gut sample per microcolony. In addition, three workers from each source colony were dissected to determine variation in microbiome composition among source colonies. DNA for these bee samples and for samples of fresh pollen balls was extracted using the Qiagen DNeasy PowerSoil kit, following included protocol, but with modified overnight incubation [41].

From these samples, we assessed fungal abundance and examined fungal and bacterial species composition. To assess fungal abundance, we used SYBR-based qPCR to determine fungal copy number in each crop and gut sample. Samples were run on a Bio-Rad CFX96 Thermal Cycler, using the ITS86F (5’ – GTGAATCATCGAATCTTTGAA – 3’) [42] and ITS4 (5’ – TCCTCCGCTTATTGATATGC – 3’) [43] primers. Run conditions were as follows: 2 minutes at 95°C followed by 40 cycles of 15 seconds at 95°C, 30 seconds at 59°C, and 30 seconds at 60°C. Fluorescence at 520nm was measured at the end of each cycle. All samples were run in triplicate, with internal standards included in each plate. To convert from the relative Cq scores produced from qPCR to total ITS copy number, the internal standard included in all qPCR plates was run through a standard PCR using the same primers and conditions as in the qPCR. This amplified DNA was then run on an agarose gel and the ITS band was removed. From this band, DNA was extracted using a Qiaex II Gel Extraction Kit and quantified using Qubit to determine ng/uL. Total copy number was calculated from this concentration using the fragment mass. A dilution curve of this amplified DNA was then run through qPCR with the original standard to determine the conversion between Cq and total copy number.

To assess fungal and bacterial species composition, we performed amplicon sequencing of the ITS and 16S rRNA (V6/8) gene region via Illumina MiSeq, implemented at the Integrated Microbiome Resource at Dalhousie University [36]. Sequence data were processed using the package “dada2” [37] using the default pipeline parameters for bacterial and fungal samples. *B. vosnesenskii* 16S sequences had low merging success, so only forward reads were processed through the pipeline and used for downstream analysis. Following assignment of sequences to amplicon sequence variants (ASVs), taxonomy was assigned for bacterial sequences using the Silva database (v 132) [38] and for fungal sequences using a hybrid UNITE/dITSy database [39]. Sequences not annotated as bacteria or fungi were removed from the datasets prior to subsequent analysis. Several samples returned a low number of sequences (<50) and corresponding low ITS copy number values. We chose to include all of these samples in our analysis to prevent bias by low DNA concentration in some sample types and treatments. Analyses excluding these samples produced qualitatively similar results to those reported here, and more detailed results are reported in Supplemental Results. Rarefaction curves for each sample were created using “vegan” [49]. Prior to community analysis, samples were normalized by dividing counts of each ASV by total ASV counts to obtain relative abundance of each ASV within each sample. To obtain genus-level absolute abundance measures, the total copy number of each sample (determined by qPCR) was multiplied by the relative abundance of genera within that sample.

### Statistical analysis

To test the effect of fungicide and fungi treatments on microcolony survival, we used a Cox proportional hazards model (“survival” package [44]) using fungicide and fungi treatment as factors, and created survival curves using the “survminer” package [45]. We analyzed per-bee nectar consumption of microcolonies using a linear mixed effects model (package “lme4” [46]) with fungicide and fungi treatments as fixed factors and source colony as a random factor. We chose to analyze per-bee nectar consumption as microcolonies often contained different numbers of bees due to mortality, which greatly impacted nectar consumption. Because egg production was low among *B. impatiens* microcolonies, we used a hurdle model (package “pscl” [47]) to account for both the probability of egg production and egg count among fungicide and fungi treatment. Egg mass was analyzed using an ANOVA incorporating fungicide treatment and fungi treatment as factors.

To test the effects of treatments on fungal abundance (ITS copy number), we used a linear mixed effect model with fungicide treatment, fungi treatment, and organ (crop/gut) as factors, and source colony as a random effect. All fungal abundance measures were log-transformed prior to analysis.

For each bee species separately, we examined if fungicide treatment, fungi treatment, or organ affected the Shannon diversity of bacterial and fungal communities using a linear mixed effects model with source colony as a random effect.

To visualize community composition of the microbiome, we used PCoA plots of Bray-Curtis dissimilarity metrics, created using the package “phyloseq” [48]. We tested differences in species composition across treatments using PERMANOVA in the “vegan” package [49] based on Bray-Curtis dissimilarities, with fungicide and fungi treatments, organ, and source colony as fixed factors. Differences in variance across treatment groups were analyzed using a multivariate version of Levene’s test for homogeneity of variances in “vegan”, with either fungicide treatment, fungi treatment, organ, or source colony as factors. To assess if ASV relative abundance differed between fungicide or fungi treatment groups, we used “DESeq2” [50] to extract differentially abundant ASVs and report results with false discovery rate (FDR) < 0.01 (Supplemental Figure S3). All data were analyzed in R version 3.6.3 [40].

## Acknowledgements

We thank Neal Williams, John Mola, Robert Schaeffer, and Rosemary Malfi for help with bumble bee rearing protocols. We thank Marshall McMunn, Jake Francis, Amber Crowley-Gall, Shawn Christensen, Richard Karban, Adam Pepi, Eric LoPresti, Vincent Tan, and Naomi Murray for feedback on previous manuscript drafts. This work was supported by Academic Senate New Research Initiatives grant awarded to R.L.V. and D.R., George H. Vansell scholarships awarded to D.R., and Microbiome Graduate Research Award to D.R.

## Data accessibility statement

All data and scripts used in this manuscript will be available on Dryad following manuscript acceptance. All datasets and metadata have been included in submission for review. Raw sequence data is available on the NCBI Sequence Read Archive (BioProject ID PRJNA759617, https://dataview.ncbi.nlm.nih.gov/object/PRJNA759617?reviewer=p085edmnjmdaeejkp5pfab9jhq).

## Notes

https://dataview.ncbi.nlm.nih.gov/object/PRJNA759617?reviewer=p085edmnjmdaeejkp5pfab9jhq

### Competing Interest Statement

The authors have declared no competing interest.

