## Supplemental Information for "Bee-associated fungi mediate effects of fungicides on bumble bees"

### Supplemental Methods

*Y-tube assays*

Two y-tube assays were performed to test volatile preferences of *B. vosnesenskii* workers. The first of these assays compared preferences between control nectar volatiles and volatiles of nectar with 7.5 ppm propiconazole. Workers were removed from source colonies and starved for two hours (n = 24 bees). One hour into this starvation period, workers were presented with samples of each nectar solution on filter paper to allow them to taste both solutions prior to testing. Each bee was released into the y-tube under red light for a total of 10 minutes. Halfway through this assay, the treatments were flipped to opposite arms of the y-tube. The time spent in each arm of the y-tube was recorded, as was each bee's first choice.

A second preference assay was run to compare volatiles of control nectar and nectar inoculated with $1x10^6$ cells/mL of the nectar yeast *Metschnikowia reukaufii* (n = 15 bees/treatment). Before fungicide treatments were applied to microcolonies, one worker was randomly pulled from each, and the above methods were repeated. After one week of feeding on their fungicide or control diets, one worker was again randomly pulled from each microcolony, and the assay was repeated.

*Statistical analysis*

Y-tube preference data for fungicide vs. control trials was analyzed using a T-test to compare total time spent in the control arm and in the fungicide arm. Y-tube preference data for yeasts vs. control trials was analyzed using an ANOVA using differences between time spent in yeast and control arms as the response variable and fungicide treatment as a predictor.

Rarefaction curves for samples were created using the 'rarecurve' function in the 'vegan' package. To determine association between focal fungal taxa, linear models were run on log-transformed abundances (calculated from relative abundance amplicon data and ITS copy number data), with other focal taxa and total fungal abundance as predictors. Crop samples were excluded from these analyses to prevent pseudoreplication within a microcolony.

### Supplemental Results

**Y-tube foraging**

Y-tube foraging assays were carried out for *B. vosnesenskii* only. When given the choice between volatiles of control nectar and volatiles of nectar inoculated with the yeast *Metschnikowia reukaufii*, workers spent more time in the arm of the y-tube containing control volatiles than the arm containing yeast volatiles, regardless of fungicide treatment ($F_{1, 202} = 12.94$, $p < 0.001$). When given a choice between control nectar volatiles and nectar volatiles with fungicide, bees showed no preference ($F_{1,40} = 1.21$, $p = 0.28$)

**Fungal microbiome**

From two pollen samples, we recovered 23,326 sequences, which were clustered into 69 ASVs. 54% of sequences belonged to Basidiomycota, and the other 46% belonged to Ascomycota. At the family level, 52% of sequences belonged to Melampsoraceae, 13% to Ascosphaeraceae, and 12% to Mycosphaerellaceae.

*Associations between focal taxa*

In *B. vosnesenskii*, there was a weak positive association between *Ascosphaera* abundance and *Starmerella* abundance ($F_{1,28} = 4.59$, $p = 0.041$), but no relationship between *Ascosphaera* and *Zygosaccharomyces* abundance ($F_{1,28} = 0.12$, $p = 0.73$). Additionally, there was no association between *Zygosaccharomyces* abundance and *Starmerella* abundance ($F_{1,29} = 0.14$, $p = 0.72$).

In *B. impatiens*, there was a positive association between *Ascosphaera* abundance and *Starmerella* abundance ($F_{1,27} = 8.55$, $p = 0.007$), but no association with *Zygosaccharomyces* ($F_{1,27} = 1.96$, $p = 0.17$) or *Debaryomyces* abundance ($F_{1,27} = 0.002$, $p = 0.97$). There was no association between *Starmerella* and *Zygosaccharomyces* abundance ($F_{1,29} = 0.79$, $p = 0.38$), between *Starmerella* and *Debaryomyces* abundance ($F_{1,29} = 0.56$, $p = 0.46$), or between *Zygosaccharomyces* and *Debaryomyces* abundance ($F_{1,29} = 1.61$, $p = 0.21$).

*Differentially abundant taxa by treatment*

Using DESeq2, we found eight fungal ASVs that differed in relative abundance between *B. vosnesenskii* microcolony treatment groups (Fig S3A). In *B. impatiens* we found three fungal ASVs that differed between treatment groups (Fig S3B).

*Analyses excluding low-read* B. impatiens *samples*

Four samples with less than 50 recovered sequences were removed from the dataset prior to diversity and community composition analyses. Fungicide and fungi treatment interacted to impact Shannon diversity ($F_{1,55} = 4.22$, $p = 0.045$), with fungi addition increasing fungal diversity, particularly in microcolonies that had previously been exposed to fungicide. Crop samples hosted more diverse fungal communities than gut samples ($F_{1,55}$ = 11.14, p = 0.002).

Fungal community composition differed between fungi treatments ($F_{1,54}$ = 4.11, p = 0.006), but not between fungicide treatments ($F_{1,54}$ = 1.52, p = 0.16), or their interaction ($F_{1,54}$ = 1.64, p = 0.12). Communities were similar between crop and gut samples ($F_{1,54}$ = 1.56, p = 0.15), and between source colonies ($F_{1,54}$ = 1.11, p = 0.32). Variance was homogenous across fungicide treatments ($F_{1,58}$ = 0.13, p = 0.72), fungi treatments ($F_{1,58}$ = 0.019, p = 0.89), crop and gut samples ($F_{1,58}$ = 0.003, p = 0.96), and source colonies ($F_{1,58}$ = 0.054, p = 0.82).

Using PCoA axes as predictors of survival, we found that neither axis 1 ($\chi^2$ = 0.47, df = 1, p = 0.49) nor axis 2 ($\chi^2$ = 0.45, df = 1, p = 0.50) was associated with differences in the survival of *B. impatiens* microcolonies.

**Bacterial microbiome**

A total of 1,546,471 sequences were obtained from 64 *B. vosnesenskii* samples, ranging from 12 to 90,730 per sample (24,163 ± 2,857, mean ± SE). Sequence recovery was not impacted by fungicide treatment ($F_{1,18}$ = 0.015, p = 0.90), fungi treatment ($F_{1,18}$ = 0.028, p = 0.87), or the interaction between these treatments ($F_{1,18}$ = 0.18, p = 0.68). Source colonies yielded different numbers of reads ($F_{7,18}$ = 3.21, p = 0.022), and sequences were recovered in much lower numbers from crop samples compared to gut samples ($F_{1,18}$ = 9.39, p = 0.0067). Sequences were clustered into a total of 891 ASVs using the dada2 pipeline. All samples reached saturation (Figure S1C). The phylum Firmicutes dominated samples, accounting for 94.7% of sequences, with Proteobacteria accounting for another 3.6% of sequences. At the family level, 73% of sequences were assigned to Carnobacteriaceae and 12.8% to Lactobacillaceae.

A total of 376,641 sequences were obtained from 64 *B. impatiens* samples, ranging from 0 to 33,552 per sample (5,885 ± 124, mean ± SE). Sequence recovery was not impacted by fungicide treatment ($F_{1,48}$ = 0.14, p = 0.71), fungi treatment ($F_{1,48}$ = 0.021, p = 0.89), or the interaction between these treatments ($F_{1,48}$ = 0.046, p = 0.83). Most sequences were recovered from gut samples, while very few were recovered from crop samples ($F_{1,48}$ = 61.25, p < 0.001). Similar number of sequences were recovered from each source colony ($F_{1,48}$ = 2.22, p = 0.14). Sequences were clustered into a total of 702 ASVs. All samples reached saturation (Figure S1D). Proteobacteria made up 81% of sequences with Firmicutes accounting for another 15% of sequences. At the family level, 67% of sequences were assigned to Neisseriacaea, 14% to Orbaceae, with smaller portions of sequences assigned to Lactobacillaceae, Bacillaceae, and Bifidobacteriaceae.

*Alpha diversity*

In *B. vosnesenskii*, we found no difference in bacterial Shannon diversity between fungicide treatment ($F_{1,55}$ = 0.50, p = 0.48), fungi treatment ($F_{1,58}$ = 2.06, p = 0.16), nor the interaction between these treatments ($F_{1, 56}$ = 1.56, p = 0.22). Diversity was also similar between crop and gut samples ($F_{1,52}$ = 0.029, p = 0.87).

For *B. impatiens* microcolonies, bacterial Shannon diversity decreased with fungicide treatment, but only in crop samples (fungicide treatment x organ interaction, $F_{1,50}$ = 5.97, p = 0.02). Fungi treatment ($F_{1,50}$ = 0.65, p = 0.43) and the interaction between fungicide and fungi treatments ($F_{1,50}$ = 0.043, p = 0.84) had no impact on diversity. Gut samples hosted more diversity than crop samples ($F_{1,50}$ = 285, p < 0.001).

*Microbiome composition*

For *B. vosnesenskii* microcolonies, the composition of bacterial communities was affected by the interaction between fungicide and fungi treatments (PERMANOVA, $F_{1,18}$ = 2.46, $R^2$ = 0.041, p = 0.019, Fig S5A). Communities also varied by source colony, but not significantly ($F_{7,18}$ = 1.51, $R^2$ = 0.18, p = 0.077). Communities were similar between crop and gut samples ($F_{1,18}$ = 0.94, p = 0.44). The variance of fungicide treatments (betadisper, $F_{1,62}$ = 1.22, p = 0.27), fungi treatments ($F_{1,62}$ = 1.06, p = 0.29), organ type ($F_{1,62}$ = 0.18, p = 0.68), and source colonies ($F_{7,56}$ = 2.01, p = 0.064) were similar. We isolated three ASVs using DESeq2 that differed in relative abundance between treatment groups. Two Bacillaceae ASVs were found in all treatments except the group receiving fungicide and fungi. A Lactobacillus ASV was found only in the group receiving fungicide and fungi.

For *B. impatiens* microcolonies, fungi treatment impacted bacteria community composition, though not significantly ($F_{1,42}$ = 1.69,  $R^2$ = 0.019. p = 0.054, Fig S5B). There was no effect of fungicide treatment ($F_{1,42}$ = 1.57, p = 0.097) or the interaction of fungicide and fungi treatments on community composition ($F_{1,42}$ = 1.56, p = 0.091). Communities varied between crop and gut samples ($F_{1,42}$ = 14.1, $R^2$ = 0.15, p = 0.001), between source colonies ($F_{1,42}$ = 10.66, $R^2$ = 0.12, p = 0.001), and was also dependent on the interaction between these two terms ($F_{1,42}$ = 8.59, $R^2$ = 0.094, p = 0.001). There was no difference in community variance by fungicide treatment ($F_{1,56}$ = 0.55, p = 0.49), fungi treatment ($F_{1,56}$ = 0.36, p = 0.53), or source colony ($F_{1,56}$ = 0.13, p = 0.72). Variance did differ between crop and gut samples, with crop samples much more tightly clustered than gut samples ($F_{1,56}$ = 23.17, p < 0.001). Using DESeq2, we found two different ASVs between microcolony treatment groups. One Bacillaceae ASV was detected mainly in control samples, and one Carnobacteriaceae ASV was present mostly in the fungi only treatment.

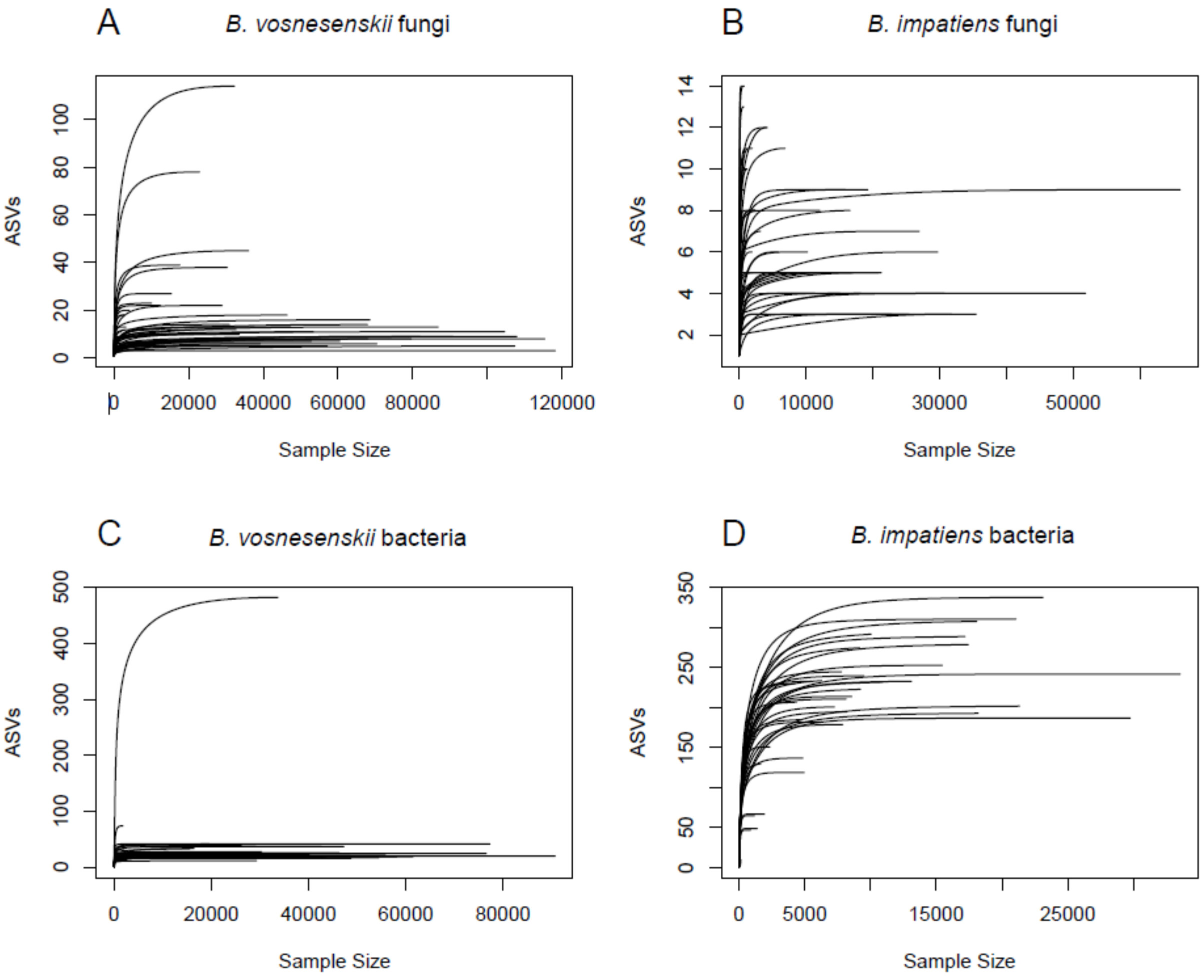


**Figure S1**. Rarefaction curves for A) *B. vosnesenskii* fungal ASVs, B) *B. impatiens* fungal ASVs, C) *B. vosnesenskii* bacterial ASVs, and D) *B. impatiens* bacterial ASVs.

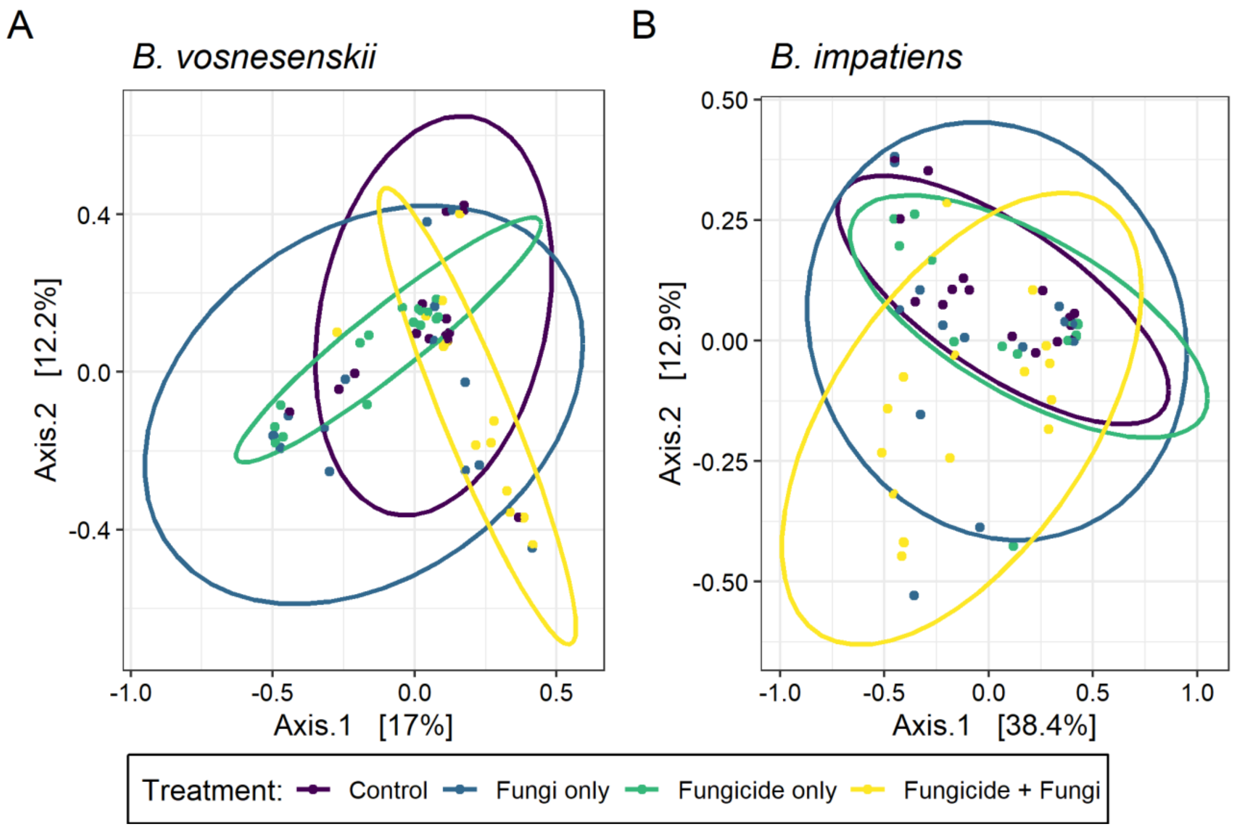


**Figure S2.** PCoA plots of fungal communities of (A) *B. vosnesenskii* and (B) *B. impatiens* microcolonies, based on Bray-Curtis dissimilarity metrics. Community composition of *B. vosnesenskii* microcolonies differed between fungicide and microbe treatments ($F_{1, 60} = 3.67$, $R^2 = 0.054$, $p = 0.001$). Community composition of *B. impatiens* microcolonies differed by microbe treatment ($F_{1,60} = 3.72$, $R^2 = 0.056$, $p = 0.007$). N = 8 microcolonies per treatment per species.

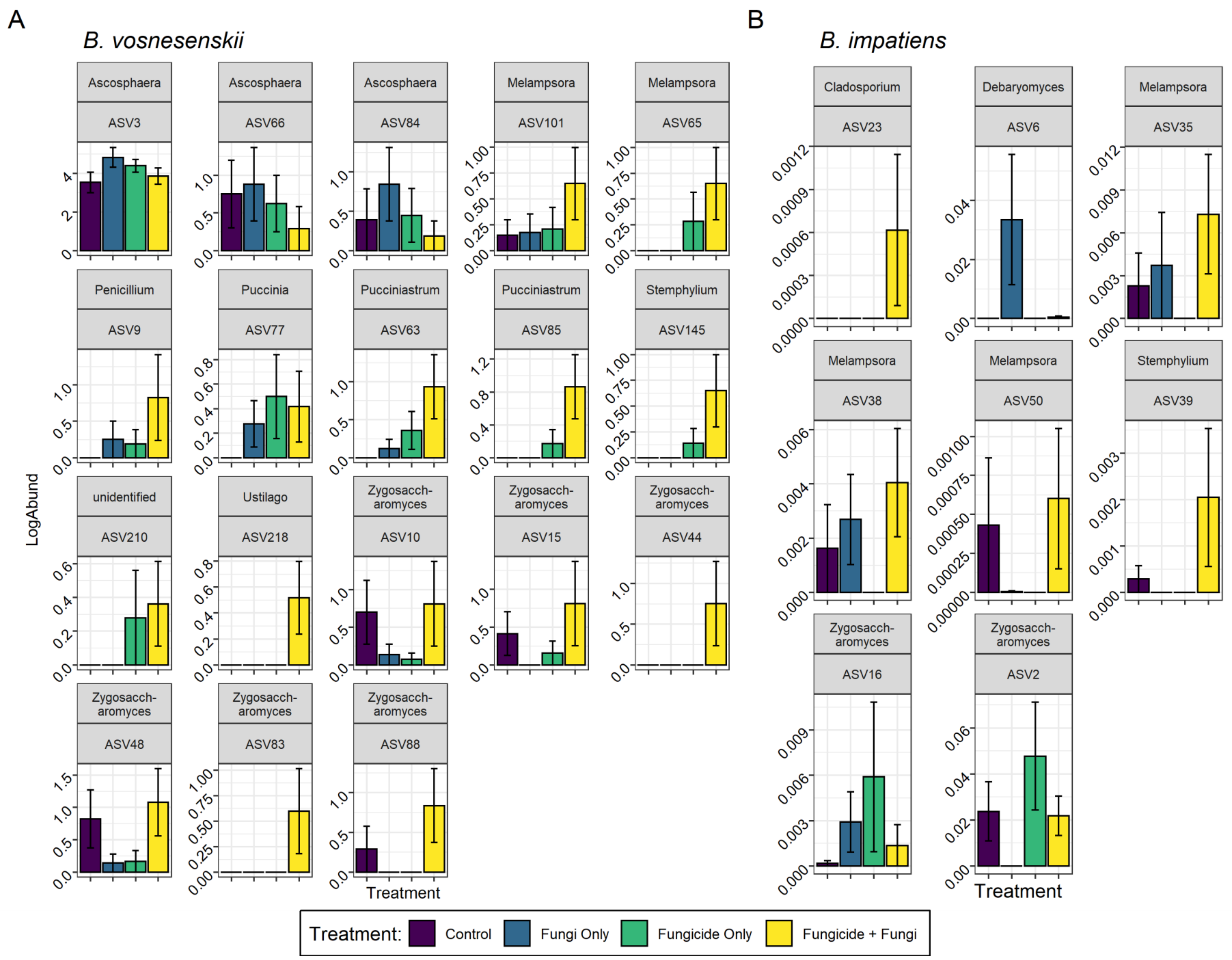
A
B. vosnesenskii
B
B. impatiens
LogAbund
Ascosphaera ASV3
Ascosphaera ASV66
Ascosphaera ASV84
Melampsora ASV101
Melampsora ASV65
Penicillium ASV9
Puccinia ASV77
Pucciniastrum ASV63
Pucciniastrum ASV85
Stemphylium ASV145
unidentified ASV210
Ustilago ASV218
Zygosacch-aromyces ASV10
Zygosacch-aromyces ASV15
Zygosacch-aromyces ASV44
Zygosacch-aromyces ASV48
Zygosacch-aromyces ASV83
Zygosacch-aromyces ASV88
Cladosporium ASV23
Debaryomyces ASV6
Melampsora ASV35
Melampsora ASV38
Melampsora ASV50
Stemphylium ASV39
Zygosacch-aromyces ASV16
Zygosacch-aromyces ASV2
Treatment
Treatment
Treatment: Control Fungi Only Fungicide Only Fungicide + Fungi

**Figure S3.** Differentially abundant fungal ASVs across fungicide and fungi treatments of (A) *B. vosnesenskii* and (B) *B. impatiens* microcolonies according to tests of differential expression in the "DESeq2" package. The log-transformed relative abundance of each ASV is shown on the y-axis, using different scales for each ASV. N = 8 microcolonies per treatment per species.

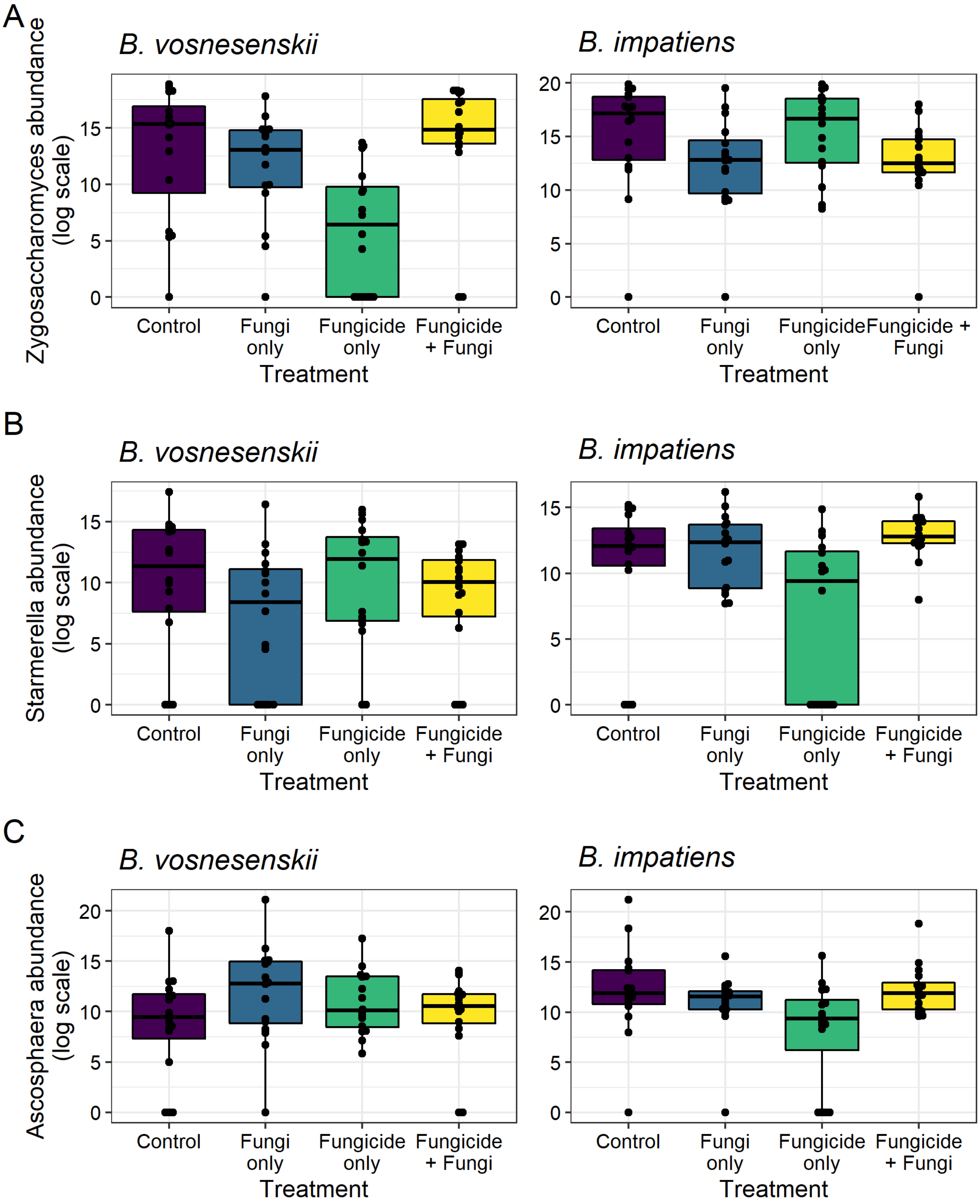


**Figure S4.** Abundance of specific fungal groups A) *Zygosaccharomyces*, B) *Starmerella*, and C) *Ascosphaera* represented by log-transformed ITS copy number. The left column represents *B. vosnesenskii* samples and the right column represents *B. impatiens* samples. N = 8 microcolonies per treatment per species. In *B. vosnesenskii*, *Zygosaccharomyces* abundance was lowest in the fungicide-only treatment ($F_{1,56} = 11.33$, $p = 0.001$), while in *B. impatiens* it was lower in fungi-treated microcolonies ($F_{1, 55} = 6.79$, $p = 0.012$). *Starmerella* abundance was not impacted by treatment in *B. vosnesenskii*, but was lowest in the fungicide-only treatment in *B. impatiens* ($F_{1,55} = 5.12$, $p = 0.028$). *Ascosphaera* abundance was highest in the fungi-only treatment in *B. vosnesenskii* ($F_{1,54} = 14.00$, $p < 0.001$), and was lowest in the fungicide-only treatment in *B. impatiens* ($F_{1,56} = 7.35$, $p = 0.009$).

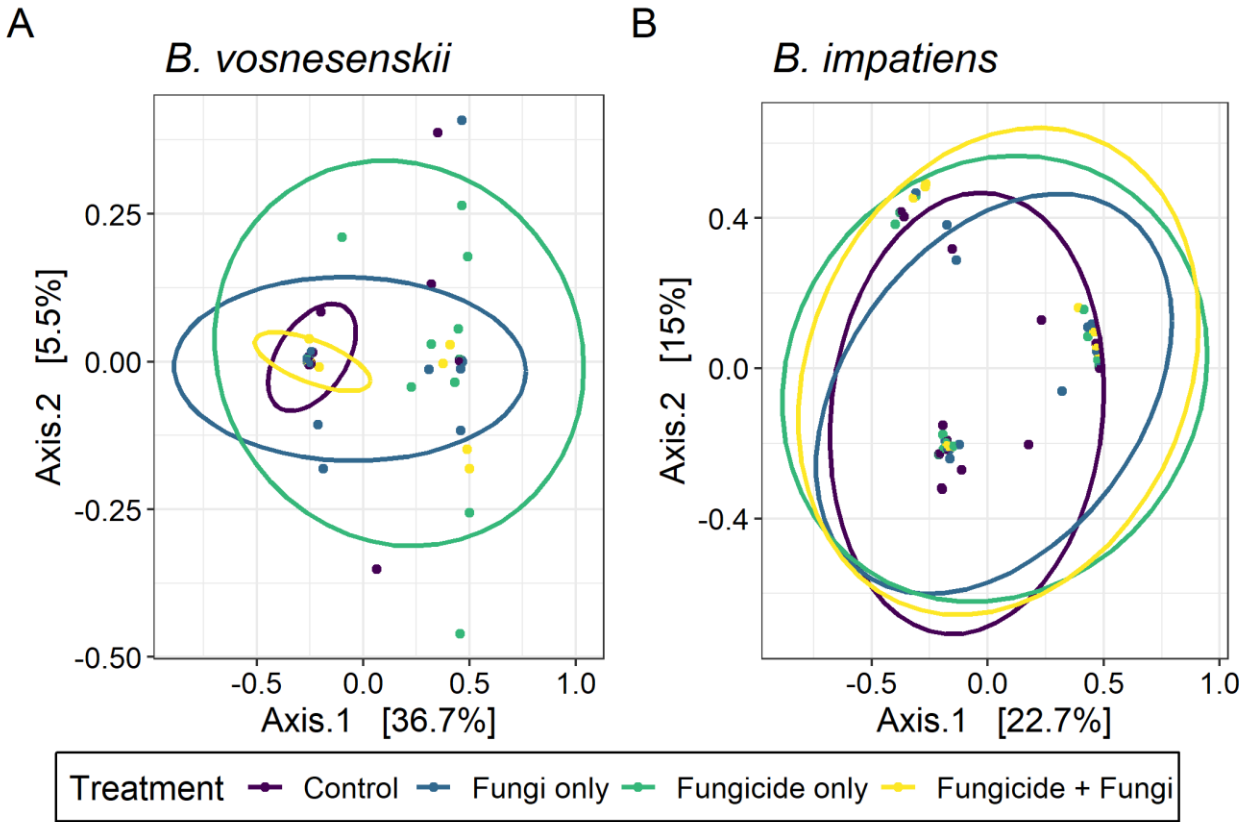


**Figure S5**. Bacterial community composition of A) *B. vosnesenskii* and B) *B. impatiens* microcolonies. Fungicide and fungi treatments interacted to impact composition in *B. vosnesenskii* ($F_{1,18} = 2.46$, $R^2 = 0.041$, $p = 0.019$). There was a marginally significant effect of fungi treatment on *B. impatiens* community composition ($F_{1,42} = 1.69$, $R^2 = 0.019$. $p = 0.054$).
